# Alpha-synuclein alters the faecal viromes of rats in a gut-initiated model of Parkinson’s disease

**DOI:** 10.1101/2021.03.29.437468

**Authors:** S. R. Stockdale, L. A. Draper, S. M. O’Donovan, W. Barton, O. O’Sullivan, L. A. Volpicelli-Daley, A. M. Sullivan, C. O’Neill, C. Hill

## Abstract

Parkinson’s disease (PD) is a chronic neurological disorder associated with the misfolding of alpha-synuclein (α-syn) into Lewy body aggregates within nerve cells that contribute to their neurodegeneration. Recent evidence suggests α-syn aggregation may begin in the gut and travel to the brain along the vagus nerve, with microbes a potential trigger initiating the misfolding of α-syn. However, changes in the gut virome in response to α-syn alterations have not been investigated. In this study, we show longitudinal changes in the faecal virome of rats administered either monomeric or preformed fibrils (PFF) of α-syn directly into their enteric nervous system. Differential changes in rat viromes were observed when comparing monomeric and PFF α-syn. The virome β-diversity changes after α-syn treatment were compounded by the addition of LPS as an adjunct. Changes in the diversity of rat faecal viromes were observed after one month and did not resolve within the study’s five month observational period. Overall, these results suggest that microbiome alterations associated with PD may, partially, be reactive to host α-syn associated changes.

## Introduction

Parkinson’s disease (PD) is the second most common progressive neurodegenerative disorder, affecting 0.1-0.2% of the population at any given time. This frequency increases with age, with more than 1% of individuals over 60 living with PD ^1^. While the overall cost of PD varies from country to country, in the UK conservative and liberal estimates calculated the annual cost of the illness between £449 million and £3.3 billion, respectively ^2^. As the mean age of populations increase, the social and economic burden of PD will rise. In fact, worldwide cases of PD are expected to double to 14 million by 2040 ^3^; therefore, a greater understanding of the disease and new interventions are urgently required.

Genetic and epidemiological studies investigating PD have not demonstrated a specific aetiology that triggers PD. Instead, many hypothesise that PD is a multifactorial disease involving a complex combination of host genetics and environmental factors ^4^. The pathophysiological onset of PD is associated with the loss of dopamine-producing neurons in the pars compacta sub-region of the substantia nigra structure of the midbrain ^5^. However, patients typically only display Parkinsonism, the motor features associated with PD, after 50-80% of dopaminergic neurons are lost ^6^. The presynaptic neuronal protein alpha-synuclein (α-syn) is linked to the loss of dopaminergic neurons, whereby the monomeric form of α-syn aggregates within nerve cells into toxic Lewy body (LB) formations that disrupt cellular homeostasis ^7^.

Gastrointestinal (GI) comorbidities are frequently associated with PD, with cohorts from independent studies reporting that 70-78% of patients experience excessive saliva, 30-97% of individuals have difficulty swallowing, and 20-89% report constipation ^8^. The non-motor symptoms associated with PD may also involve α-syn, as its aggregation is detected earlier and more frequently in the enteric (i.e. gastrointestinal) nervous system of PD patients ^9,10^. Recent animal models have also demonstrated the spread of pathologic α-syn along the gut-brain axis through the vagus nerve ^11–13^. This has led researchers to investigate the GI tract and its community of microbes as a potential environmental trigger for the aggregation of α-syn that may contribute to the development of PD.

There are 10^13^ to 10^14^ bacteria, archaea, fungi, and viruses associated with the human body, known collectively as the microbiota. Non-culture-based approaches to study their genomic material (microbiome) have demonstrated the GI composition of PD patients differs from controls ^14–21^. In addition, transplantation of human faecal microbiotas from either PD patients and non-PD controls to mice resulted in a motor function decline amongst the mice humanized with a PD microbiota ^22^. Specific bacterial taxa, and even bacterial-infecting viruses (termed bacteriophages or phages), have been proposed as potential contributors to PD ^17,23^. However, caution is often required when interpreting microbial changes associated with specific conditions, as alterations observed in the microbiome may be causal, or reactive, to human host-associated changes ^24^.

In this study, we investigated if the viral component of the rat microbiome (virome) changes in response to α-syn injected into rat enteric nervous systems. This study was specifically performed in young rats to distinguish if α-syn could be associated with virome changes independent of microbial changes that would occur during PD development in older rats. Both the monomeric and aggregated fibrillar forms of α-syn were injected directly into the myenteric plexus of surgically-prepared rat duodenal walls, with and without intraperitoneal injection of bacterial lipopolysaccharide (LPS) as a potential inflammatory adjunct. Virome diversity changes were observed between rats receiving specific α-syn treatments, and also longitudinally across individual rats. Specific viruses and VCs were noted as differentially abundant across treatment groups, with microbiome changes reacting to α-syn and potentially PD-inducing conditions.

## Results

### Defining the rat faecal virome

Viruses of the rat microbiome were identified using database-dependent and -independent approaches, applied to both the viral-enriched and whole-genome sequencing (WGS) data (see Methods). Viral sequences identified in the viral-enriched and WGS data were merged into a single database and made non-redundant by removing duplicate sequences and retaining the larger of any two homologous sequences (≥ 90% identity and ≥ 90% coverage). The majority of sequences in the final rat viral database originated from the viral-enriched fraction of the microbiome (3,506/4,009 sequences). The largest viral contig was 291,209bp in length, while the mean and median length of viral contigs is 4,123 and 1,621bp, respectively (Figure 1a).

**Figure 1.**
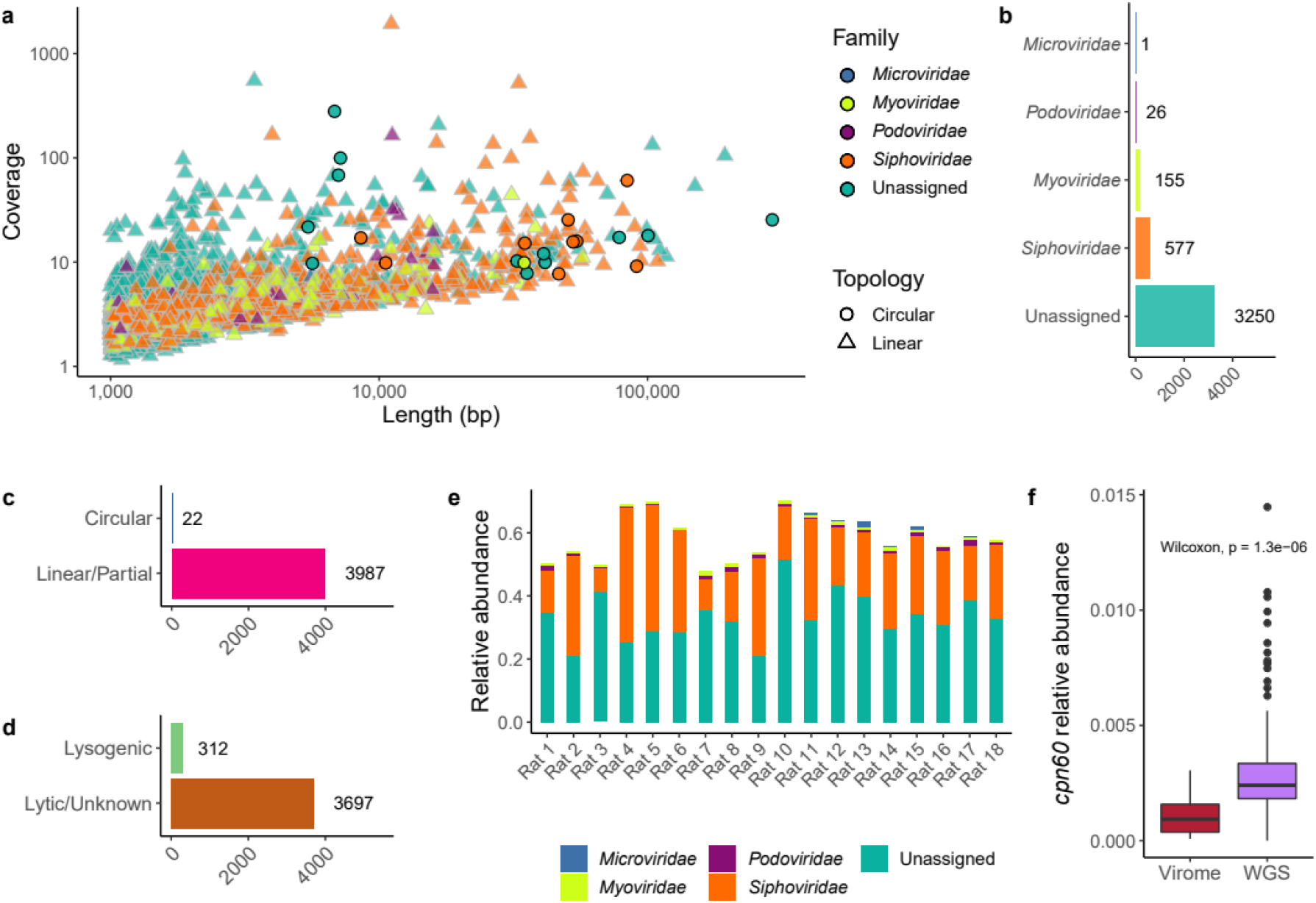
The rat faecal virome. **(a)** Contig length (bp) versus sequencing coverage of identifiable viruses present in the faeces of male rats. The shape aesthetic was used to highlight the topology of assembled viral contigs, while colour represents proposed taxonomic assignment. **(b)** The breakdown of rat faecal viruses into their putative familial-level taxonomic assignments. **(c)** The topology of identified viral contigs, whereby circular sequences are most likely complete genomes. **(d)** Viral contigs encoding a phage integrase and/or recombinase were annotated as lysogenic phages. **(e)** The relative abundance of viral contigs within the virome sequencing data, aggregated by taxonomic assignment. Sequencing reads aligning to contigs not recognised as viral are omitted. **(f)** The relative abundance of the bacterial house-keeping gene *cpn60* detected in the virome sequencing data versus the WGS data.

The viral nucleic acid extraction and sequencing protocols used in this study did not employ random DNA amplification, such as multiple displacement amplification (MDA), known to bias the viral composition towards small circular genetic elements. Therefore, the relative abundance of specific viral taxa such as small single-stranded DNA *Microviridae* may be more accurate than previous faecal virome studies ^25^. The average raw sequencing reads per sample for rat faecal viromes was 2,926,309 (max. 3,616,794, min. 2,221,657). The rat faecal viromes analysed in this study were dominated by bacterial-infecting phage. However, the term virome (and not “phageome”) is used throughout this study, as this study’s viral-detecting protocol does not preclude the identification of rat-infecting viruses, and they may be present amongst the ‘Unclassified’ viruses.

Of the tailed *Caudovirales* phages assigned a putative familial taxonomic rank, *Siphoviridae* phages were the most abundant, with *Myoviridae* and *Podoviridae* phages detected at decreasing frequencies (Figure 1b). All viral sequences identified within this study were clustered into pseudogenera termed viral clusters (VCs) using the vContact2 gene sharing network-based algorithm. A total of 920 VCs were generated, with the largest cluster containing 20 sequences.

The majority of phage contigs identified in this study are linear (99.45%; Figure 1c). While the rat faecal virome might have relatively few circular genomes, it is more likely that greater sequencing depth is required to complete and circularize more viral genome sequences. The viral sequences identified in this study were queried for the presence of phage integrases and/or recombinases. Only 8.4% of phages possessed genes encoding for lysogenic conversion that would facilitate integration and replication as a prophage (Figure 1d).

Using a database-dependent and -independent approach to identify viruses, the average relative abundance of the identified rat faecal viruses accounts for 58.9% of the virome sequencing reads generated (Figure 1e; min. 47.6%, max. 70.2%). Therefore, a significant proportion of the rat faecal virome is not analysed and could be considered as “viral dark matter” ^26^. While significantly lower than within the whole shotgun sequencing, there are instances where the bacterial housekeeping gene, *cpn60*, is detected within the virome sequencing data (Figure 1f). Therefore, potential bacterial contamination may have been inadvertently categorised as viral, and a greater understanding of the rat microbiome will be required to distinguish the origin of sequences.

### The rat microbiome changes over time

The male Sprague-Dawley rats employed in this study were approx. two months of age at the study’s onset and seven months old at its conclusion. Therefore, the potential impact of time in shaping the rat faecal microbiome was investigated. During the MetaPhlAn2 analysis of the WGS data, an average of 99.7 and 0.3% of the microbiome sequencing reads that could be assigned to microbial taxa were classified as bacterial and viral, respectively. A longitudinal axis to the rat faecal virome was obtained by aligning the WGS sequencing data onto the final rat viral database. An average of 44,431 WGS sequencing reads aligned to viral sequences (min. 572; max. 256,910; median 33,298).

There was a statistically significant change in the faecal virome and total microbiome composition of the rats over the study’s duration, with time explaining 14.6% and 13.3% of their variances, respectively (Figures 2a & b). The two-dimensional separation of faecal samples by dissimilarity (β-diversity) was not driven by intra-sample composition (α-diversity). No statistical difference in α-diversities after Bonferroni correction are observed in rat faecal viromes by treatment group, examining either virome or WGS data (Supplementary Table 1). The PCoA axes 1 and 2 for the microbiome data explain more of the variation than for the virome data (28.5 and 15.5% versus 21.1 and 9.4%, respectively).

**Figure 2.**
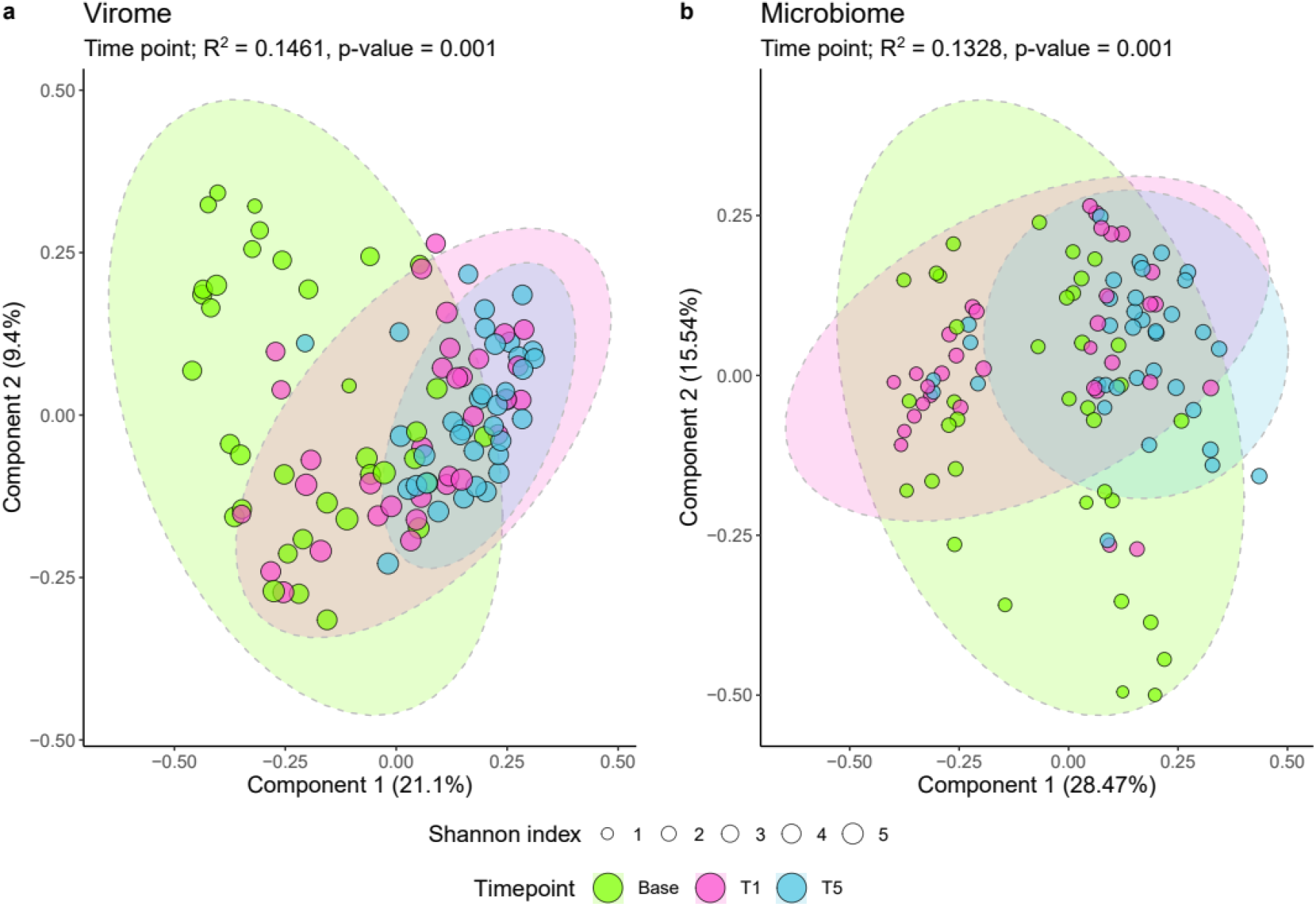
α- and β-diversity analysis of the rat faecal microbiome. Study time points T1 and T5 represent one and five months, respectively, and are grouped by ellipses. Two dimensional PCoA ordination of rat faecal samples, using the Bray-Curtis dissimilarity index, demonstrates the β-diversity between samples. The size of the sample points analysed represents their α-diversity Shannon index values. **(a)** WGS sequencing reads mapped onto the identified rat faecal virome, and **(b)** relative abundance of bacterial species in the WGS data.

### α-syn alters the rat faecal virome

As it is established that the rat faecal virome changed over the course of the study, any alterations occurring in α-syn injected rats need to be examined in light of the control treatments, Sham and LPS. The Sham and LPS controls experienced 22.6 and 19.5% variance, respectively, in their viromes over the study’s duration (Figures 3a & b). Less than a 1% difference in the variance of the rat faecal viral compositions are observed in the rats that received the α-syn monomer or α-syn monomer with LPS (28.0 and 27.6% variances explained, respectively; Figures 3c & d). However, the effect of the α-syn monomer on rat viromes is far greater than differences attributable to either the Sham or LPS control treatments. The PFF form of α-syn appears to have had only a small effect on the rat virome relative to the Sham and LPS controls; however, the PFF plus LPS treatments caused the greatest virome alteration (variances explained 23.9% and 30.8%, respectively; Figures 3e & f).

**Figure 3.**
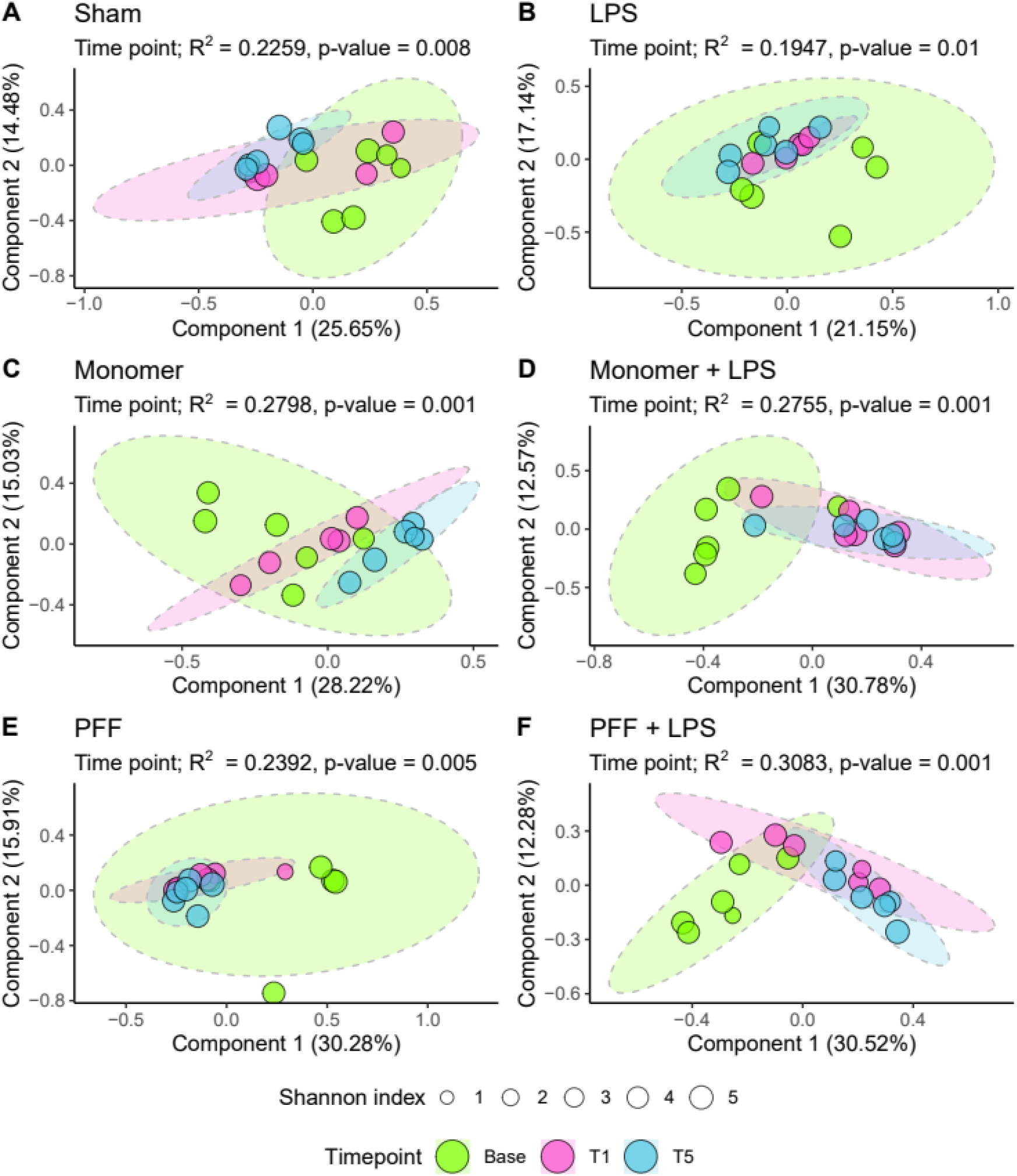
α- and β-diversity analysis of the rat faecal virome by treatment condition. Ellipses group samples by the study time points investigated. The size of sample points represents their Shannon index α-diversity, while the two dimensional PCoA ordination of sample points represents their β-diversity dissimilarity using the Bray-Curtis index. The treatment conditions investigated in this study are; **(a)** Sham, **(b)** LPS, **(c)** α-syn monomer, **(d)** α-syn monomer and LPS, **(e)** α-syn PFF, and **(f)** α-syn PFF and LPS.

### Changes in individual rat viromes

In order to investigate if changes in the rat faecal virome were directional, whereby a sample is not merely more or less similar to its baseline through random fluctuations, each individual rat’s faecal virome was investigated relative to its baseline state. Using Bray-Curtis index values that indicate the dissimilarity of two given virome samples, it was observed that the greatest variation in the rat faecal virome occurs between Base and T1, compared to T1 and T5 (Figure 4a). There was no statistical difference in the dissimilarity values between the ‘Base vs T1’ and ‘Base vs T5’. Additionally, the virome at five months was more frequently dissimilar to the baseline compared to the one-month time point (Figure 4b).

**Figure 4.**
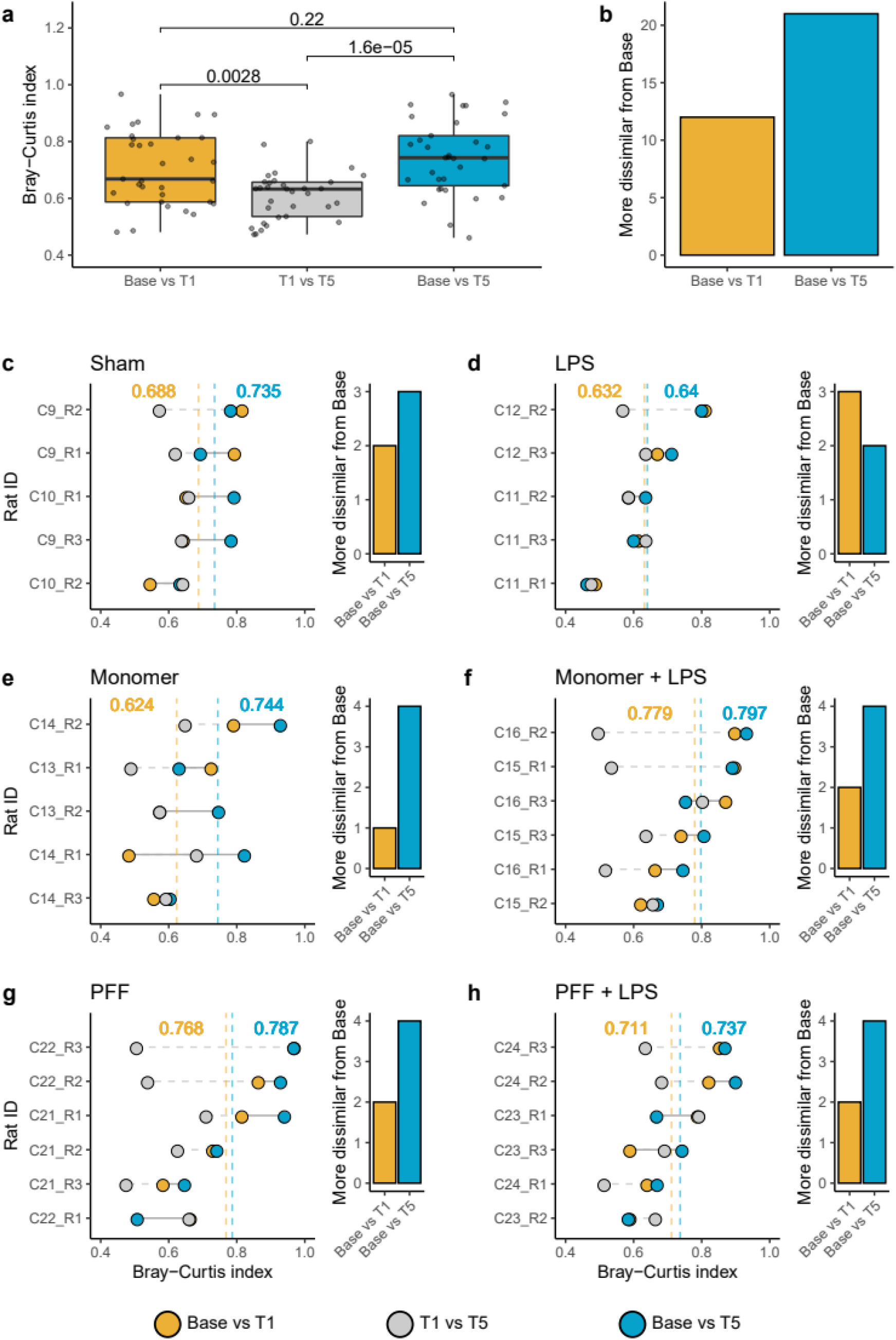
Changes in the faecal virome of individual rats, separated by the treatment conditions. **(a)** The Bray-Curtis dissimilarity index values for all rat faecal viromes between study time points, with Wilcoxon test p-values shown. Points represent the specific Bray-Curtis index values. **(b)** Summary of whether faecal viromes analysed were more dissimilar from the study’s baseline (Base) using the Bray-Curtis index after 1 month (T1) or 5 months (T5). The left-hand image of treatment panels **(c-h)** demonstrates the Bray-Curtis dissimilarity for individual rat faecal viromes. A horizontal solid grey line connects the ‘Base vs T1’ and ‘Base vs T5’ Bray-Curtis values, while a horizontal dashed grey line indicates the Bray-Curtis change between T1 and T5. Vertical dashed lines and numerical insets are colour coded to indicate the average Bray-Curtis index values at the respective time points. The right-hand image of each treatment panel specifies if T1 or T5 Bray-Curtis values are more dissimilar from the Base.

For the individual rat viromes segregated by treatment conditions (Figures 4c-h), the least variation was observed between the average Sham, LPS, and α-syn monomer treatment baselines versus T1 time points (Figures 4c-e). Whereas within the first month, the α-syn monomer plus LPS, the PFF α-syn, and the PFF α-syn plus LPS treatments had the least similarity when averaged across rats (Figures 4f-h). The viromes of rats receiving the α-syn monomer plus LPS are the most dissimilar to their baseline after the study’s five months (Figure 4f).

Additionally, we investigated if virome changes associated with specific α-syn treatments were the result of cage effects. Each rat was assigned a numerical value and a code based on its cage. For example, code C21-R2 specifies cage number 21 and rat 2. Two cages, with 2-3 rats per cage, were used for each treatment. Looking at each individual rat, no specific trend across any of the treatment conditions was observed, such as all the rats within a specific cage dramatically differing at T1 or T5 relative to their baseline (Figures 4c-h). Also, the average Bray-Curtis dissimilarity values were similar for each specific treatment across the two corresponding cages.

### Viral changes caused by α-syn treatments

If the α-syn treatments investigated in this study altered the rat microbiome and/or virome, it is expected that the abundance of viruses would be significantly altered in one or more time points (p-value < 1E-05). The Sham and LPS treatment arms of this study resulted in the fewest differential viral and VC changes (Figure 5a & b). There were only two viruses increased at the one-and five-month time points of the Sham-administered rats, while no viruses were consistently altered amongst LPS treated rats.

**Figure 5.**
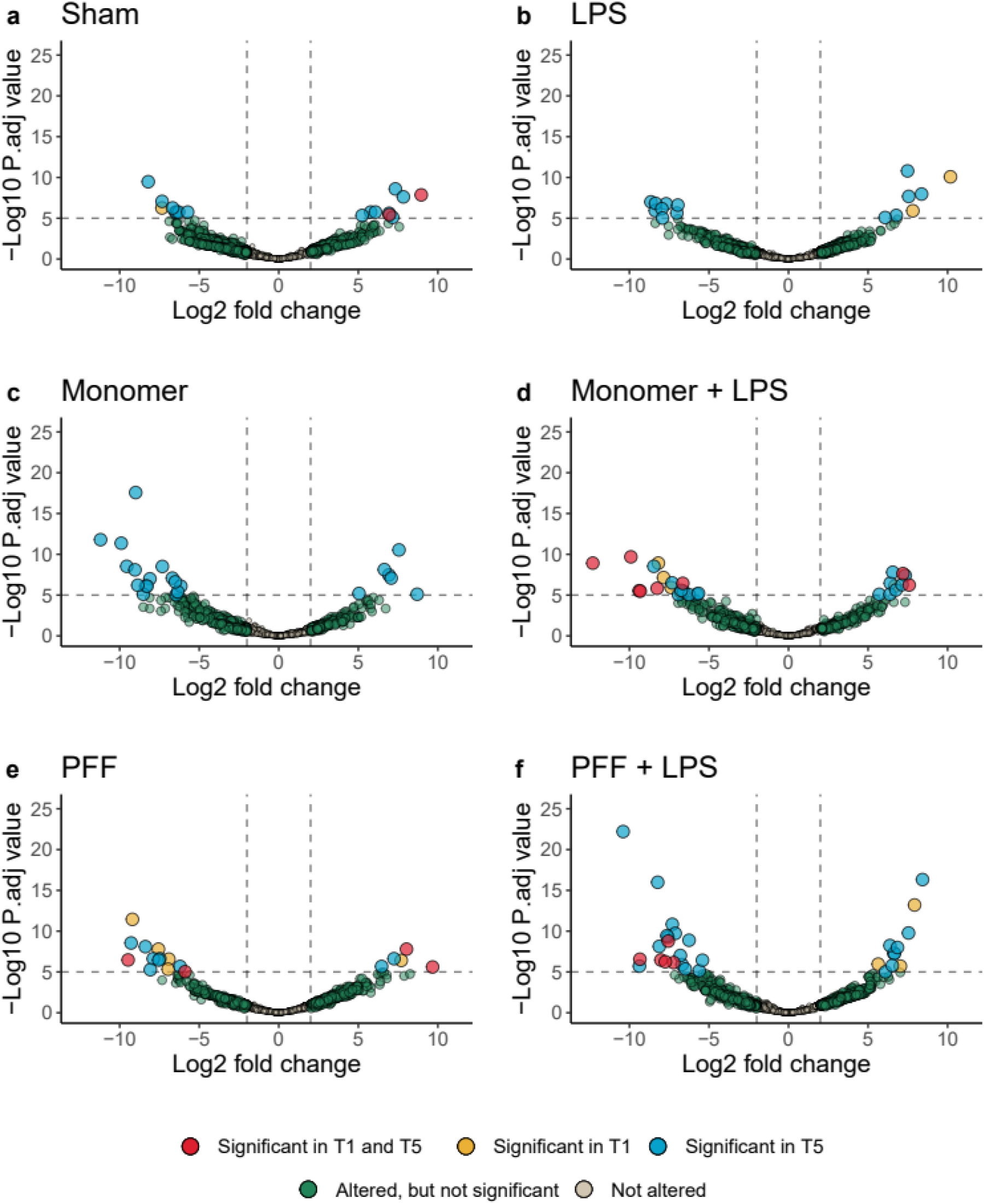
Volcano plots showing differential abundance of rat faecal viruses and viral clusters at one month (T1) and five months (T5) relative to the study’s baseline. The vertical dashed lines of each panel indicate log2 fold changes equivalent to +2 or -2, while the horizontal dashed line denotes a Bonferroni adjusted p-value significance of 1E-05. The axis values of all panels are scaled equivalently.

For rats receiving the monomeric form of α-syn, 16 and 6 viruses had decreased and increased differential abundance at the five-month time point (Figure 5c). The α-syn monomer treatment did not induce a change in a virus or virus cluster across the time-points investigated. The α-syn monomer plus LPS, the PFF, and the PFF plus LPS treatment arms of the study all induced significant abundance changes in viruses or VCs consistent across T1 and T5 time points. These treatments caused decreased differential abundance in six, two, and five viruses and viral clusters, respectively (Figures 5d-f). While the α-syn monomer plus LPS and the PFF treatments both increased two viral groups across both time points (Figures 5d & e).

### Common viral changes across PD inducing treatments

In order to discern if a common pattern of changes occurs across different treatment conditions, all viruses and viral clusters that were noted to have a differential abundance at any time point were compared across the treatments tested. A total of 78 viruses and viral clusters, predicted from the virome sequencing and the WGS sequencing data, were recorded as differentially abundant at one or more time points relative to their baseline (Figure 6). Of these, the majority (48 of 78, 61.5%) were only altered in one treatment condition. These are most probably the result of random changes that have occurred in the rat microbiome over the course of the study.

**Figure 6.**
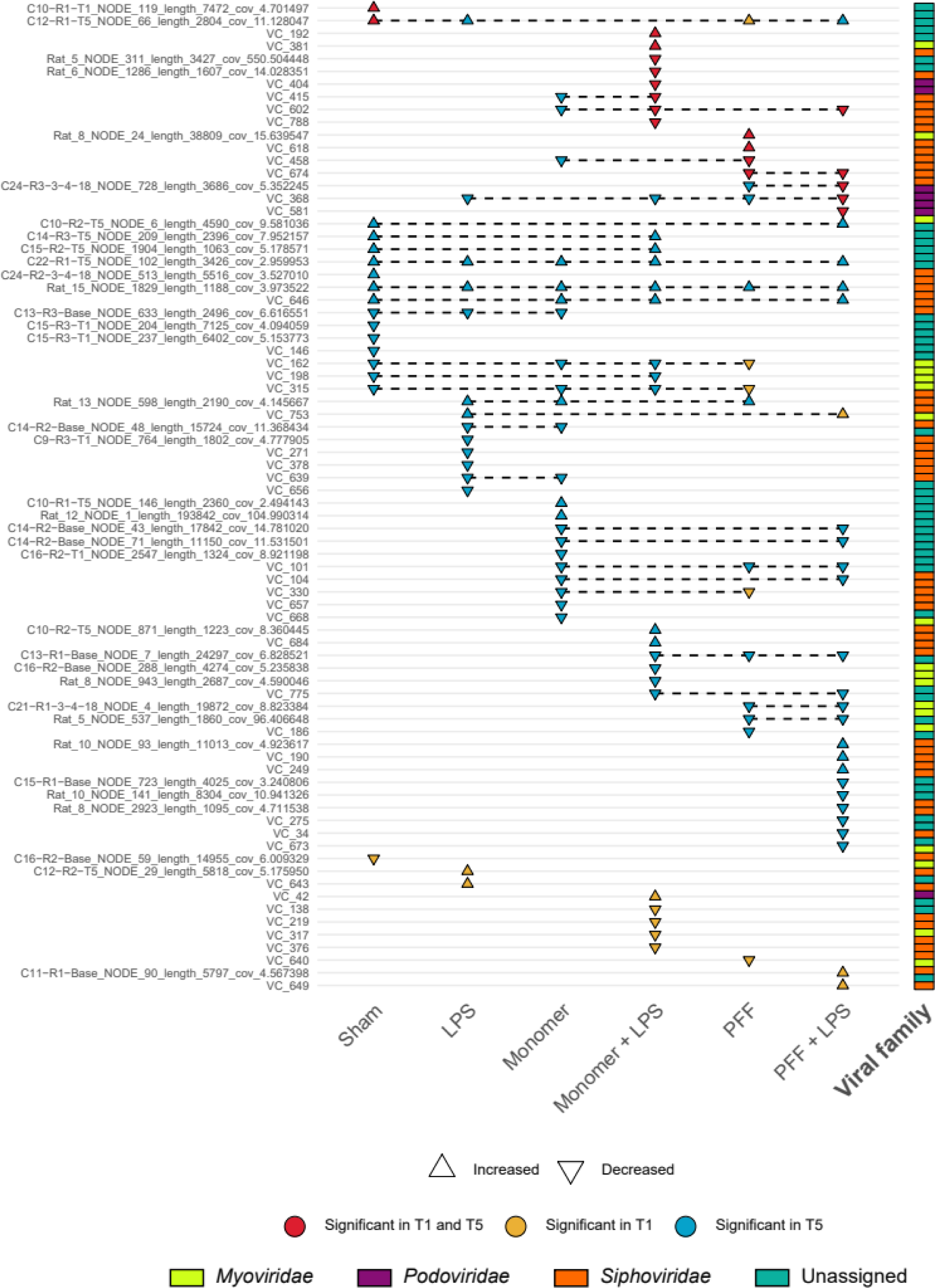
Viruses and viral clusters which were differentially abundant at statistical significance in at least one of the study’s time points, relative to the baseline. Viruses or viral clusters that were differentially abundant across two or more treatment groups are connected using black dashed lines. The shape of each point indicates if the virus or viral cluster had an increased or decreased differential abundance. The putative familial taxonomic assignment of the various viruses is indicated to the right-hand side of the image.

Nine viruses or VCs were altered across both the T1 and T5 time points analysed, supporting the hypothesis that these are permanent microbiome changes. Only four viruses are concordantly changed across the Sham and LPS controls, seven in the α-syn monomer and α-syn monomer plus LPS treatments, and finally, nine in the α-syn PFF and α-syn PFF plus LPS treated rats. There was one virus with an increased differential abundance at T5 across all treatment groups (contig Rat_15_NODE_1829_length_1188_cov_3.973522). Whereas a second viral contig (C22−R1−T5_NODE_102_length_3426_cov_2.959953) has an increased differential abundance detected in all but the α-syn PFF treatment group. Therefore, these two contigs may be fragments of the same altered virus, with the latter sequence putative annotated as a *Siphoviridae* phage.

## Discussion

There are a limited number of viral metagenomic analyses conducted on wild rats as they are a potential source of zoonotic infections ^27–29^. However, despite their frequent use as models of human diseases, to our knowledge, the virome of lab rats has not been investigated (see Methods). In this study, the majority of faecal viruses analysed are putatively annotated as tailed phages belonging to the *Siphoviridae, Myoviridae*, and *Podoviridae* families, with few *Microviridae* observed. A similar trend is observed in the accumulative relative abundance of viral taxa within microbiome sequencing data (Supplementary figure 1). These top-level hierarchical viral taxa are similarly observed in the faeces of mice and humans ^30,31^.

Unlike many mammalian faecal virome analyses, advances in sequencing technologies have enabled us to conduct our analysis without a nucleic acid amplification step. Therefore, the relative abundance of viral taxa and overall diversity of viromes presented here is likely to be more accurate than previous studies. With a sequencing depth of approximately 3M reads per sample, fewer genomes than expected were circular. Without the random amplification bias towards small circular viral elements, and an overall better representation of the virome diversity within the microbiome, an even greater sequencing depth may be required to complete metagenomic viral genomes. On average, there were more lytic than lysogenic phages noted in the rat virome, and this did not statistically differ over the study’s duration (Supplementary figure 2). However, the relative abundance of lytic phages decreased (p-value < 0.05) in the monomeric α-syn plus LPS and PFF treatment groups, relative to the study’s Sham treatment group. This was investigated because changes in the ratio of lytic to lysogenic phages were observed in a comparison of healthy and inflammatory bowel disease human viromes ^32^.

The majority of the rat faecal virome can be considered as ‘dark matter’, whereby assembled sequences are unlike those deposited in reference databases. This is similar to studies of the human virome, where 40-90% of sequences were unidentifiable ^33^. In this study, an average of 58.9% of the rat virome sequencing data could be aligned back to the viral database generated from the sequencing of their faecal viromes. Therefore, while a proportion of the virome sequencing data was not analysed, the remaining uncharacterised data was a cautionary measure to prevent unintentional interpretation of bacterial-contaminating sequences as viral.

The diversity of rat faecal viromes changed over the study’s five-month duration. As we are not aware of any published studies on the longitudinal rat faecal virome, it is difficult to ascertain the cause(s) for this variation. Interestingly, the magnitude of change observed for the whole microbiome and virome was similar using the same methodological procedures (variances of 14.6% and 13.3%, respectively). While this result may be coincidental, it would be logical that changes in bacterial populations would have a concomitant effect on the abundance of their viruses. One potential explanation for the shifts observed in the microbiome and virome of rats over five months would be their accelerated aging relative to humans. When comparing their lifespans, every 13.8 rat days approx. equates to a human year ^34^. Therefore, rats were potentially first analysed in this study during puberty and reached social maturity at its conclusion. The virome changes associated with these developmental periods are poorly described across all mammalian hosts.

Knowing that the rat faecal virome changes over time, all fluctuations associated with inter-sample diversity must be interpreted cautiously. The variance (R^2^) explained by viral β-diversities did not differ dramatically between the Sham, LPS, or α-syn PFF treatments (22.6%, 19.5%, and 23.9%, respectively). However, large β-diversity differences occurred over the study’s duration between control groups and rats administered the α-syn monomer, α-syn monomer plus LPS, and α-syn PFF plus LPS treatments (28.0%, 27.6%, and 30.8%, respectively). Therefore, specific α-syn treatments affected the overall longitudinal β-diversity of rats differently, with the fibril form of α-syn with LPS injection as an inflammatory adjunct resulting in the greater variance.

When broken down into an investigation of individual rat viromes, the greatest variation was observed between the baseline and the first month, T1. However, the viromes of rats continued to change between T1 and T5, and more frequently the viromes were more dissimilar from their basal state at the five-month time point, T5. The magnitude by which individual rat viromes transformed over the study was smallest for the Sham, LPS, and α-syn monomer treatments. Once more, the greatest changes observed in individual rats occurred in those receiving the α-syn monomer plus LPS, the α-syn PFF, and α-syn PFF plus LPS treatments.

The individual viruses and VCs responsible for the rat faecal virome changes were investigated. A similar trend to the β-diversity analysis was observed across all of the study’s treatment groups. More viruses were differentially altered after five months, compared to one month. Amongst the Sham rats investigated, only two viruses of VCs were differentially altered at one month with the change maintained until the five-month time point. The LPS treated control rats had no virus or VC altered across both time points. The abundance and magnitude of viral changes are amplified in rats treated with the α-syn monomer plus LPS, the α-syn PFF, and the α-syn PFF plus LPS treated rats.

Changes in rat faecal viromes were examined for consistent patterns across treatment groups. Ideally, specific viruses would be altered across the various α-syn treated rats indicating a consistent effect on viral composition. However, few viruses and viral clusters were differentially abundant across α-syn treatments, examining either the monomeric or fibrillar form of α-syn with and without LPS (Supplementary figure 3). Taken together, viral changes in the rat faecal virome may be indirectly associated with inflammatory or microbial changes associated with α-syn in the gut. However, no specific alteration in the virome could be definitively linked to the native or misfolded α-syn, despite several viruses differentially abundant across different treatments.

In conclusion, we present the first analysis of the lab rat faecal virome. The study focused on the effect caused by the addition of α-syn on the viral composition over five months. There is clear evidence that the virome of rats changed following the injection of α-syn into the enteric nervous system. The magnitude of change observed across faecal viromes was consistently greater amongst rats administer α-syn monomer plus LPS, α-syn PFF, or α-syn PFF plus LPS. However, additional studies are required to clearly establish if α-syn induced virome alterations are associated with the pathogenesis or gastrointestinal comorbidities of PD.

## Methods

### Generation of α-syn PFFs

Recombinant monomeric mouse α-syn was purified and used to generate α-syn PFFs, as previously described ^35^. A Pierce LAL high capacity endotoxin removal resin was used to minimize endotoxin. Endotoxin levels were 0.017 Unit/µg of protein as determined using a Pierce LAL endotoxin quantitation kit. Briefly, α-syn monomer, kindly provided by Laura Volpicelli-Daley, was thawed rapidly at 37°C in a 1.5 ml microcentrifuge tube, then centrifuged at 100,000*g* at 4°C for 60 min. The supernatant was removed and its protein concentration was determined using a Nanodrop spectrophotometer, using the mouse α-syn coefficient 7450 M^-1^cm^-1^ and molecular weight approximately 15 kDa. The supernatant was diluted in 50 mM Tris pH 7.5, 150 mM KCL to a final protein concentration of 5 mg/ml. A proportion of the solution was retained for subsequent ENS injection of α-syn monomer. The remainder was placed in a thermomixer at 37°C and shaken at 1000 r.p.m for 7 days. Following 7 days, the clear solution became turbid, which is indicative of PFF formation. The formation of PFFs was validated using the sedimentation assay and transmission electron microscopy as previously described ^35^. Immediately prior to surgery, PFFs were sonicated at 30% power on the Misonix Sonicator 3000, 1s on, 1s off for 120s total, in two 30s sessions, in order to break the fibrils into small fragments. Transmission electron micrographs showed an aggregated protein structure, with filaments that are typical of PFFs. Measurement of fibril length after sonication revealed that > 60% of the fibrils were less than 60 nm in diameter. A sedimentation assay showed that there were approximately equal amounts of α-syn PFFs in the supernatant and pellet, indicating efficient PFF formation ^35^.

### Animal housing and husbandry

Adult male Sprague-Dawley rats were purchased from Envigo, UK and maintained on a 12h:12h light:dark cycle (lights on at 08:00h) at 21 ± 2°C and humidity (30 -50%), with access to standard rat chow and water ad libitum. All experiments were conducted in accordance with the European Directive 2010/63/EU, and under a project authorization issued by the Health Products Regulatory Authority, Ireland (AE19130/P036). Animals were randomly divided into six treatment groups, as follows: (a) Sham: bovine serum albumin (BSA) injected into duodenal wall (n=12), (b) LPS: intraperitoneal (i.p.) injection of LPS (n=12), (c) monomer: α-syn monomer injected into duodenal wall (n=12), (d) monomer plus LPS: α-syn monomer injected into duodenal wall plus an immediate i.p injection of LPS (n=12), (e) PFF: PFF injected into duodenal wall (n=12), and (f) PFF plus LPS: PFF injected into duodenal wall plus an immediate i.p injection of LPS (n=12). Animals were housed three per cage, according to their treatment group. After one month, six animals from each group were culled and both gut and brain samples were retrieved for further analysis, the other six were culled at five months.

### Administration of α-syn into the duodenal wall

Animals were anaesthetised using gaseous isoflurane (4-5% to induce anaesthesia; 2-3% to maintain anaesthesia). On the day before surgery, animals were pre-treated systemically with the non-steroidal anti-inflammatory analgesic, carprofen (Rimadyl®, 1 ml/ 500 ml drinking water). Animals were maintained at a constant body temperature by the use of a heating pad. Each animal’s abdomen was shaved and sterilised using betadine. The laparotomy procedure was conducted as follows. A 2.5 cm-incision was made vertically along the abdomen using a sterile blade, then a scissors was used to cut through the muscle layer. Approximately 3 cm in length of the stomach and the duodenum was partially exposed to aid visualisation of the injection site. A Harvard automatic pump connected by tubing to a 50 µl Hamilton syringe with a 25-gauge needle was used to perform the injections. In order to validate the injection procedure and the transmission from duodenal myenteric neurons to the CNS, 2 µl of 1 % GFP-non-toxic fluorescent cholera toxin (Thermofisher, catalogue number C34775) was injected as described previously ^36,37^ into three sites in the duodenal wall. Following this validation, 2 µl of either mouse α-syn monomer (5 µg/µl) or mouse PFF (5 µg/µl) was injected into each of these three sites. Animals in the LPS groups also received an i.p. injection of LPS (2.5 mg/kg, in 0.9 % w/v sterile saline) immediately after the laparotomy (LPS: Sigma, catalogue number L2630, 0111:B4) ^38^. Sham animals received an injection of BSA (2 µl per site, 5 µg/µl) into the duodenum, as previously described ^12,39–41^ and LPS animals only received 2.5 mg/kg LPS as i.p injection. All animals were treated for two days following surgery with the non-steroidal anti-inflammatory analgesic, carprofen (Rimadyl®, 1 ml/500 ml drinking water) to ensure pain relief.

### Faecal collection and virome shotgun sequencing

Faecal pellets were directly extracted from animals at three time points; at the pre-surgical baseline, termed: baseline, one month post-surgery (T1) and five months post-surgery (T5). Samples were stored on dry ice and then frozen at -80°C. Nucleic corresponding to the viromes of each group of rats was purified from faecal samples, with approximately half a pellet per rat included in the extraction. Faecal samples were homogenised in 10 ml SM buffer followed by centrifugation twice at 5,000g at 10°C for 10 mins and filtration through a 0.45 µm syringe filter to remove particulates and bacterial cells. NaCl (0.5 M final concentration; Sigma) and 10% w/v polyethylene glycol (PEG-8000; Sigma) were added to the resulting filtrate and incubated at 4°C overnight. Following centrifugation at 5,000g at 4°C for 20 mins, the pellet was resuspended in 400 µl SM buffer. An equal volume of chloroform (Fisher) was added and following 30 sec of vortexing the sample was centrifuged at 2,500g for 5 mins at RT. The aqueous top layer was retained and subjected to RNase I (10 U final concentration; Ambion) and DNase (20 U final concentration; TURBO DNA-free™ Kit, Invitrogen) treatment in accordance with the manufacturer’s guidelines.

To isolate the nucleic acids, virus like particles were incubated with 20 μL of 10% SDS and 2 μL of proteinase K (Sigma, 20 mg/mL) for 20 min at 56°C, prior to lysis by the addition of 100 µL of Phage Lysis Buffer (4.5 M guanidine thiocyanate; 45 mM sodium citrate; 250 mM sodium lauroyl sarcosinate; 562.5 mM β-mercaptoethanol; pH 7.0) with incubation at 65°C for 10 min. Viral nucleic acid was purified by two treatments with an equal volume of phenol:chloroform:isoamyl alcohol (25:24:1), finally passing it through a QIAGEN Blood and Tissue Purification Kit and eluting samples in 30 µL of AE Buffer. In order to include any RNA viruses in the downstream analysis, 22 µl of the eluent underwent reverse transcriptase (RT) treatment (Superscript IV, Thermo Fisher) as per the manufacturer’s instructions, albeit doubling all volumes to increase the volume of template used. Prior to library preparation this RT product was sonicated following adjustment of the volume to 52.5 µl with low-EDTA TE buffer. Shearing of unamplified DNA/cDNA mixture was performed on M220 Focused-Ultrasonicator (Covaris) with the following settings: peak power of 50 W, duty factor of 20%, 200 cycles per burst, total duration of 35 sec. All steps were performed in accordance with the manufacturer’s protocol.

In order to concentrate the DNA/cDNA fragments entering the library, it was passed through the Zymo Genomic DNA Clean & Concentrator™ Kit, eluting in 17 µl DNA Elution Buffer supplied in the kit. Library preparation was carried out using Accel-NGS 1S Plus kit (Swift Biosciences) according to manufacturer’s instructions, with the addition of a final purification step using a 1:1 DNA/AMPure beads ratio to eliminate any small fragment contamination prior to sequencing. A single-indexed pooled library was sequenced using 2×150 nt paired-end sequencing run on an Illumina Novaseq platform at Genewiz, Germany.

### Virome data processing

Illumina sequencing reads were downloaded and processed as follows. Read quality was assessed pre- and post-processing using FastQC version 0.11.3 ^42^. Adaptors and sequenced nucleotides demonstrating low quality were pruned using Trimmomatic version 0.36 in paired-end mode, when the quality dropped below a Phred score of 30 for a 4bp sliding window ^43^. While processed reads less than 70bp were dumped, the surviving paired and unpaired reads were used in all subsequent analyses. The level of bacterial contamination within the virome sequencing data was estimated by aligning reads against a *cpn60* database^44^.

Reads were assembled into contigs using metaSPAdes version 3.11.1. All contigs less than 1,000bp were discarded. Within the virome sequencing, viruses were predicted through several different procedures. The assembled sequences were queried against the NCBI viral RefSeq database version 95 (E-value 1E-05) using BLAST version 2.6.0+ ^45^. All circular contigs detected in the virome sequencing were considered as viral. Assembled nucleotide sequences were run through the VirSorter pipeline, where category 1 and 2 viruses were examined further. Viral dark matter was detected by a BLASTn search against NCBI’s NT database, with assembled sequences from the viral-enriched shotgun data with no representative in the NT database included in downstream analyses. Viral proteins were predicted using Prodigal version 2.6.3 ^46^ with the ‘meta’ option enabled and Shine-Dalgarno training bypassed. Contig-encoded proteins were queried against the Prokaryotic Viral Orthologous Groups database (pVOGs) using HMMER version 3.1.b2 ^47^. The following cut-offs were employed to detect sequences rich in viral proteins: contigs < 5kb needed ≥ 3 pVOG hits; ≥ 5 and < 10kb, 4 pVOGs; ≥ 10 and < 20kb, 5 pVOGs; ≥ 20 and < 40kb, 6 pVOGs; ≥ 40 and < 60kb, 7 pVOGs; and ≥ 60kb, 8 pVOGs. Sequences identified through the different approaches were pooled together and made non-redundant, keeping the larger of two sequences when the BLAST identity and coverage between sequences exceeded 90%.

Subsequent to the detection of putative viruses in the rat faecal virome sequencing data, potential contaminant sequences were removed as follows. The encoded proteins of all putative viruses were queried against a database of ribosomal proteins. All contigs encoding ribosomal proteins were removed from further analysis. Additionally, proteins were queried for similarity to the Pfam plasmid replication proteins: PF01051, PF01446, PF01719, PF04796, PF05732, and PF06970. Contigs containing a HMMER hit with a score ≥ 15 were discarded.

Virome sequencing reads were aligned to the final viral contigs database using the WGS sequencing data (kindly supplied by Teagasc, Moorepark). While the virome sequencing data was generated on pooled rat faecal samples, the WGS sequencing was performed on multiple time points. Reads were aligned using Bowtie2 version 2.3.4.1 in end-to-end mode ^48^. The breadth of coverage of reads mapping across each viral contig was calculated using the BEDTools version 2.26.0 coverage function ^49^. In order to determine if a viral sequence was truly present, and not the result of multiple reads stacking onto a single conserved element, the following breadth of coverage cut-offs were applied: minimum 3 sequencing reads; for contigs < 5kb, 50% of the sequence needed to be covered in aligned reads; contigs ≥ 5 and < 20kb, 30% of the sequence needed to be covered, and for contigs ≥ 20kb, 10% of the sequence needs to be covered. If the breadth of coverage filter was not met for a specific contig within a sample, zero reads were recorded.

Viral nucleotide sequences were grouped into Viral Clusters (VCs), a pseudo-genera taxonomic rank, using vContact2 version 0.9.8 ^50^. VCs were generated using default settings with the inclusion of known viruses from the Bacterial and Archaeal Viral RefSeq version 85, with ICTV and NCBI taxonomy. The sequencing reads of viruses belonging to specific VCs were aggregated. Viruses grouped into ambiguous vContact2 clusters (i.e. outliers, overlap) were treated as singletons. Viruses not grouping into VCs were nonetheless retained during the analysis. The putative familial-level taxonomic composition of viral sequences were predicted using the Demovir script (https://github.com/feargalr/Demovir) with default settings. Integrases and recombinases were identified amongst viral-encoded proteins using a probabilistic hidden Markov model (HMM) generated from Pfam protein families; PF00239, PF00589, PF02899, PF07508, PF09003, PF09299, PF10136, PF13356, PFF14659, PF16795, and PF18644.

### Virome analysis

Analyses were conducted in R version 3.6.1 ^51^, implemented through R studio. Images were drawn using ‘ggplot2’ ^52^, utilising the ‘RColorBrewer’ and ‘jcolors’ packages for colour schemes ^53,54^. Statistical comparisons were performed using the ‘ggpubr’ package ^55^, with two and multiple group comparisons executed through Wilcoxon and Kruskal-Wallis tests, respectively. α- and β-diversities of samples were calculated with the ‘vegan’ and ‘phyloseq’ packages, respectively ^56,57^. Differences in sample β-diversities displayed in this study were calculated using the Bray-Curtis dissimilarity index with Principal Coordinate Analysis (PCoA) two-dimensional ordination. Permutational multivariate analysis of variance (PERMANOVA) statistical comparisons of PCoA-grouped samples was conducted with the adonis function from the vegan package. Differential abundance changes in viruses and VCs were calculated using the ‘DESeq2’ package ^58^. Reported p-values were adjusted using Bonferroni correction. The heatmap annotation bar was generated using the ‘ComplexHeatmap’ package ^59^. Literature searches were conducted in PubMed, February 2020, using the search term: “rat” AND “phage” OR “rat” AND “phage”.

## Frequent abbreviations

PD: Parkinson’s disease
α-syn: Alpha-synuclein
LB: Lewy body
GI: Gastrointestinal
LPS: Lipopolysaccharide
WGS: Whole genome shotgun
VCs: Viral clusters
PFF: Pre-formed (α-syn) fibrils

## Acknowledgements

This publication has emanated from research conducted with the financial support of Science Foundation Ireland under Grant numbers grant numbers SFI/12/RC/2273_P2 and SFI/14/SP APC/B3032.

## Author contributions

SMOD, AS and CON conceived the study. SMOD performed the animal study. AS and CON supervised the animal study. LAD extracted and sequenced the faecal viromes. SRS performed the analysis. WB and OOS conducted the WGS sequencing. LVD purified monomeric α-syn. SRS, LAD, SMOD, CON, and CH interpreted the results and wrote the manuscript. CON and CH secured the funding.

## Competing interests

The authors declare no conflict of interest.

## Corresponding authors

Correspondence should be addressed to

## Data availability

The supplementary data and scripts required to generate the images and interpret the results of this study are openly available in Figshare at http://doi.org/10.6084/m9.figshare.14332985.

## Supplementary figure legends

**Supplementary figure 1.**
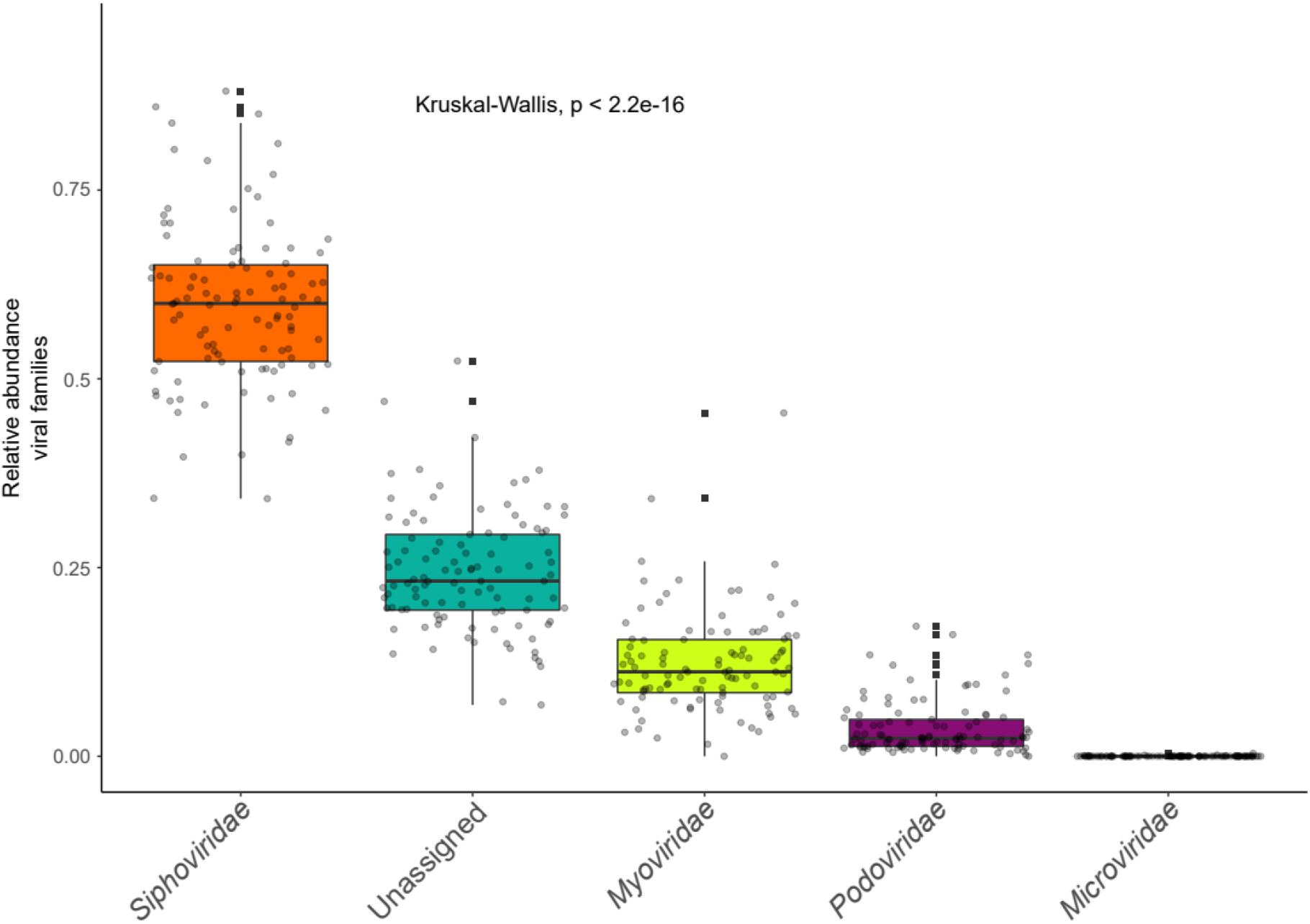
Relative abundance of predicted viral taxa in WGS data. Aggregation of sequencing data for each rat faecal sample investigated by Demovir predicted viral taxonomic assignment.

**Supplementary table 1.** Alpha-diversity analysis of rat faecal viromes. Wilcoxon-paired statistical comparisons of rat faecal virome alpha-diversities, with Bonferroni correction, using both viral and WGS sequencing data.

|  | Data | Group 1 | Group 2 | P-value | Bonferroni<br>P-value | Significant |
| --- | --- | --- | --- | --- | --- | --- |
| 1 | Virome | Sham | LPS | 0.394 | 1.000 | ns |
| 2 | WGS | Sham | LPS | 0.0177 | 0.270 | * |
| 3 | Virome | Sham | Monomer | 0.122 | 1.000 | ns |
| 4 | WGS | Sham | Monomer | 0.4745 | 1.000 | ns |
| 5 | Virome | Sham | Monomer + LPS | 0.083 | 1.000 | ns |
| 6 | WGS | Sham | Monomer + LPS | 0.2447 | 1.000 | ns |
| 7 | Virome | Sham | PFF | 0.062 | 0.920 | ns |
| 8 | WGS | Sham | PFF | 0.0380 | 0.570 | * |
| 9 | Virome | Sham | PFF + LPS | 0.525 | 1.000 | ns |
| 10 | WGS | Sham | PFF + LPS | 0.1837 | 1.000 | ns |
| 11 | Virome | LPS | Monomer | 0.357 | 1.000 | ns |
| 12 | WGS | LPS | Monomer | 0.0145 | 0.220 | * |
| 13 | Virome | LPS | Monomer + LPS | 0.083 | 1.000 | ns |
| 14 | WGS | LPS | Monomer + LPS | 0.3687 | 1.000 | ns |
| 15 | Virome | LPS | PFF | 0.219 | 1.000 | ns |
| 16 | WGS | LPS | PFF | 0.6598 | 1.000 | ns |
| 17 | Virome | LPS | PFF + LPS | 0.782 | 1.000 | ns |
| 18 | WGS | LPS | PFF + LPS | 0.3514 | 1.000 | ns |
| 19 | Virome | Monomer | Monomer + LPS | 0.483 | 1.000 | ns |
| 20 | WGS | Monomer | Monomer + LPS | 0.0773 | 1.000 | ns |
| 21 | Virome | Monomer | PFF | 0.636 | 1.000 | ns |
| 22 | WGS | Monomer | PFF | 0.0054 | 0.081 | ** |
| 23 | Virome | Monomer | PFF + LPS | 0.386 | 1.000 | ns |
| 24 | WGS | Monomer | PFF + LPS | 0.1434 | 1.000 | ns |
| 25 | Virome | Monomer + LPS | PFF | 0.791 | 1.000 | ns |
| 26 | WGS | Monomer + LPS | PFF | 0.1038 | 1.000 | ns |
| 27 | Virome | Monomer + LPS | PFF + LPS | 0.097 | 1.000 | ns |
| 28 | WGS | Monomer + LPS | PFF + LPS | 0.8391 | 1.000 | ns |
| 29 | Virome | PFF | PFF + LPS | 0.192 | 1.000 | ns |
| 30 | WGS | PFF | PFF + LPS | 0.1427 | 1.000 | ns |

**Supplementary figure 2.**
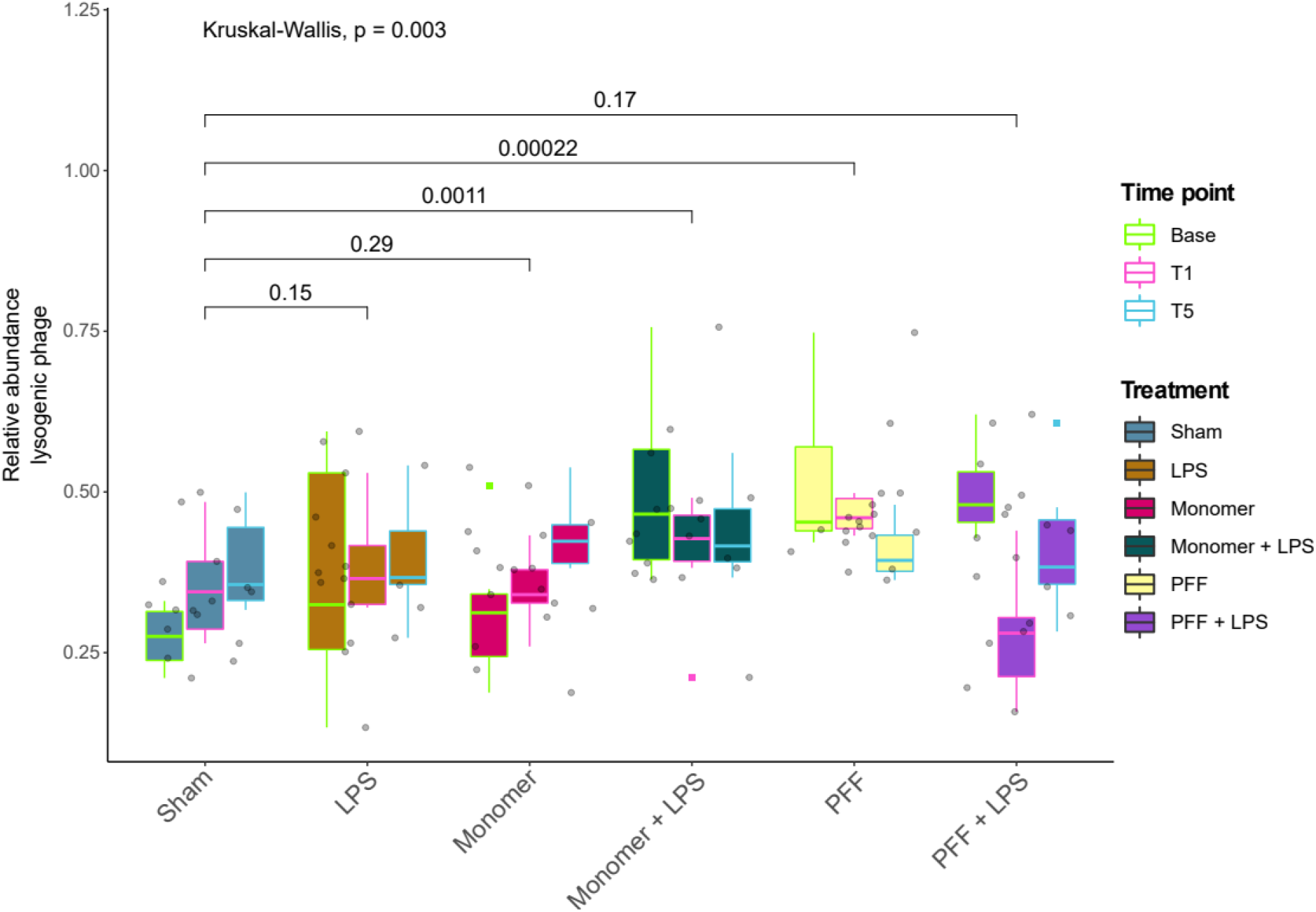
Longitudinal changes in the relative abundance of rat faecal lysogenic phages by treatment. P-values for specific group comparisons were performed using Wilcoxon-paired tests, while the Kruskal-Wallis test was performed across all groups.

**Supplementary figure 3.**
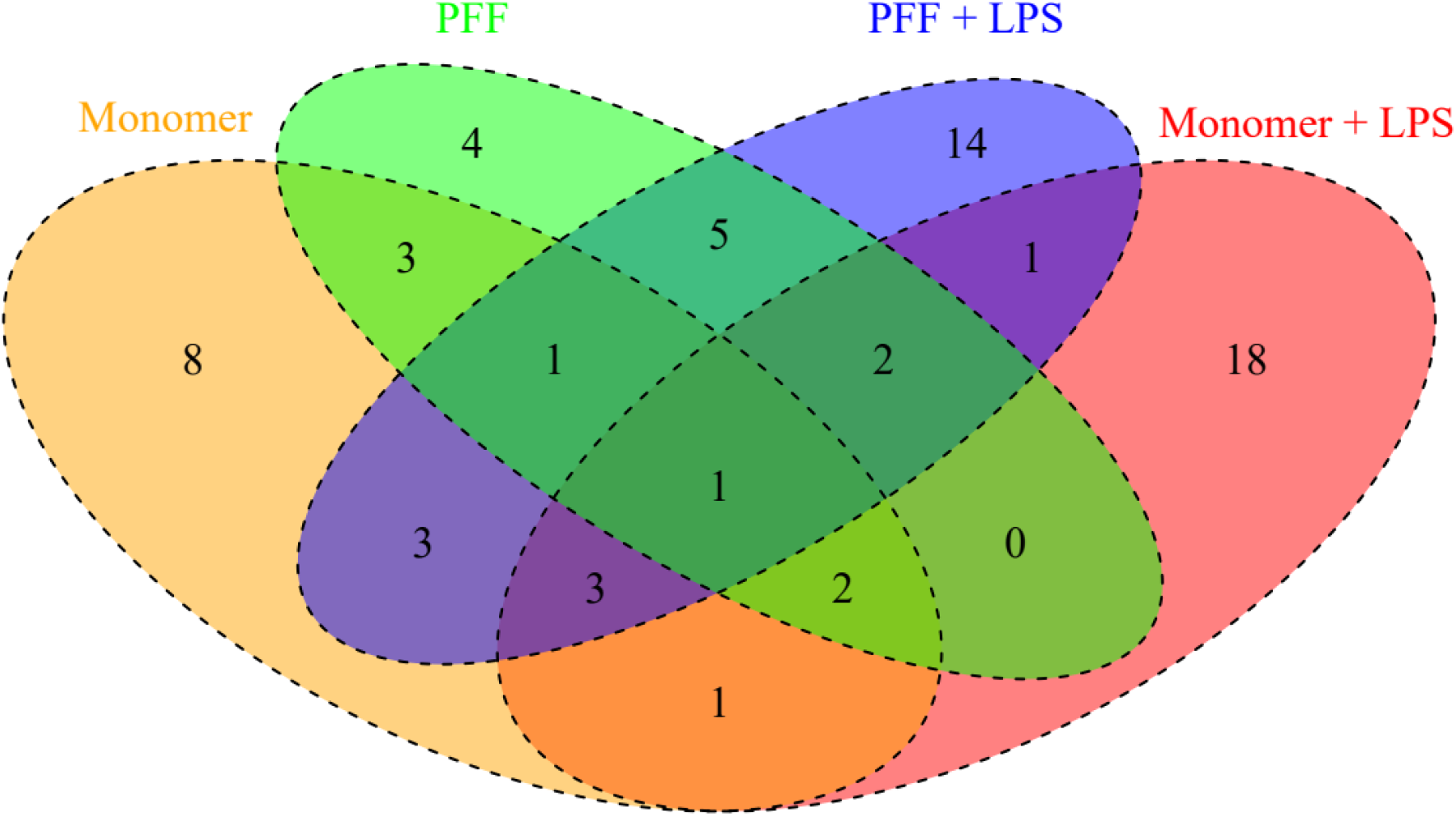
Overlapping rat faecal viruses differentially altered by alpha-synuclein treatment.

## Notes

http://doi.org/10.6084/m9.figshare.14332985

### Competing Interest Statement

The authors have declared no competing interest.

